# THE MAJOR ROLE OF JUNCTIONAL DIVERSITY IN THE HORSE ANTIBODY REPERTOIRE

**DOI:** 10.1101/2022.06.20.496904

**Authors:** Carlena Navas, Taciana Manso, Fabio Martins, Lucas Minto, Rennan Moreira, João Minozzo, Bruno Antunes, André Vale, Jonathan R. McDaniel, Gregory C. Ippolito, Liza F. Felicori

**Affiliations:** Laboratory of Synthetic Biology and Biomimetics, Departamento de Bioquímica e Imunologia, Instituto de Ciências Biológicas - ICB, Universidade Federal de Minas Gerais, Belo Horizonte, Brazil; University of Carabobo, Faculty of Health Sciences, School of Biomedical and Technological Sciences Department of Morphological and Forensic Sciences, Valencia Venezuela; The International Immunogenetics Information System / IMGT Institut de Génétique Humaine / IGH – CNRS Montpellier / France; Multi-users Laboratories Center, Instituto de Ciências Biológicas - ICB, Universidade Federal de Minas Gerais, Belo Horizonte, Brazil; Production and Research Centre of Immunobiological Products, Department of Health of the State of Paraná, Piraquara 83302-200, Brazil; Program in Immunobiology, Carlos Chagas Filho Institute of Biophysics, Federal University of Rio de Janeiro, Rio de Janeiro, Brazil; Department of Molecular Biosciences, The University of Texas at Austin, 100 E. 24th Street, Stop A5000, Austin, TX, 78712, USA

**Keywords:** horse, antibody repertoire, BCR-seq, junctional diversity

## Abstract

The sequencing of the antibody repertoire (Rep-seq) revolutionized the diversity of antigen B cell receptor studies, allowing deep and quantitative analysis to decipher the role of adaptive immunity in health and disease. Particularly, horse (*Equus caballus*) polyclonal antibodies have been produced and used since the century XIX to treat and prophylaxis of diphtheria, tuberculosis, tetanus, pneumonia, and, more recently, COVID-19. However, our knowledge about the horse B cell receptors repertories is minimal. We present a deep horse antibody heavy chain repertoire (IGH) characterization of non-immunized horses using HTS technology. In this study, we obtained a mean of 248,169 unique IgM clones and 66,141 unique IgG clones from four domestic adult horses. Rarefaction analysis showed sequence coverage was between 52 and 82% in IgM and IgG isotypes. We observed that besides horses antibody can use all of the functional IGHV genes, around 80% of their antibodies use only three IGHV gene segments, and around 55% use only one IGHJ gene segment. This limited VJ diversity seems to be compensated by the junctional diversity of these antibodies. We observed that the junctional diversity in horses antibodies is highly frequent, present in more than 90% of horse antibodies. Besides this, the length of this region seems to be higher in horse antibodies than in other species. N1 and N2 nucleotides addition range from 0 to 111 nucleotides. In addition, around 45% of the antibody clones have more than ten nucleotides in both N1 and N2 junction regions. This diversity mechanism may be one of the most important in providing variability to the equine antibody repertoire. This study provides new insights regarding horse antibody composition, diversity generation, and particularities compared to other species, such as the frequency and length of N nucleotide addition. This study also points out the urgent need to better characterize TdT in horses and in other species to better understand antibody repertoire characteristics.

## Introduction

The effective humoral immune response depends partly on having a variety of B cells with different B cell receptors (BCRs) capable of recognizing and binding to many different antigens. The entire set of B cells with different BCRs is called the antibody repertoire (Glanville et al., 2009). In humans, it has the theoretical potential to reach a size of up to 10^16^-10^18^ unique antibody sequences (Briney et al., 2019).

The sequencing of the repertoire (Rep-seq) revolutionized the antigen B-cell receptors studies, allowing deep and quantitative analysis to decipher the role of adaptive immunity in health and disease (Georgiou et al., 2014). However, besides being less common, antibody repertoire analysis in species such as chicken, sheep, pig, cattle, and horses revealed new insights into the many different mechanisms that can create antibodies diversity in vertebrates (Butler et al., 2009; Liljavirta et al., 2014; Reynaud et al., 1989; Sun et al., 2012).

Particularly, horses (*Equus caballus*) polyclonal antibodies have been produced and used since the century XIX for the treatment and prophylaxis of diseases such as diphtheria, tuberculosis, tetanus, and pneumonia (ANDERSON, 1955; Cole & Moore, 1917; Glatman-Freedman & Casadevall, 1998; Gonçalves et al., 2007; Lang et al., 2000) to the present day. It is even being used in the current COVID-19 pandemic as a treatment in some countries (Cunha et al., 2020; Zylberman et al., 2020).

Similar to other vertebrates, horses have three types of immunoglobulin chains: light lambda (IGL), light kappa (IGK), and heavy (IGH). The horse antibody V(D)J gene segments were annotated by Sun et al. (2010) and reviewed by Walter et al. (2015), using an EquCab 2.0 genome composed of several scaffolds. After that, the EquCab3.0 genome was published, and the international ImMunoGeneTics information system® (IMGT®) annotated the IG locus. In this annotation, the horse IGH locus present on chromosome 24 has 104 IGHV (21 functional, 74 pseudogenes, and nine ORFs), 44 IGHD (16 functional, twenty-eight ORFs), and nine IGHJ (six functional, and three ORFs).

So far, analyzes of the repertoire of equine antibodies have been carried out by different methodologies, most of them by Sanger sequencing, with low deepness (Almagro et al., 2006; Tallmadge et al., 2013, 2014). Only in 2019, our group carried out a deeper horse antibody repertoire analysis using the new generation technology (NGS) was carried out, showing some characteristics of 45,000 IGH clones and 30,000 IGL clones as new gene transcripts (IGHV6S1 and IGLV4S2) and the amino acids composition and features of CDR-H3 (Manso et al., 2019). However, some essential horse antibody repertoire characteristics are still unclear, such as somatic hypermutation frequency and the characteristics of the junction, among others. Furthermore, the fraction of the potential repertoire expressed in an individual is unknown, and how similar repertoires are between individuals who have lived in similar environments.

We present a deeper horse antibody heavy chain repertoire (IGH) characterization of non-immunized horses using HTS technology, where we obtained a mean of 248,169 unique IgM clones and 66,141 unique IgG clones from four domestic adult horses. Sequence coverage was between 52 and 82% in IgM and IgG isotypes. We observed that the IGHV4 subgroup is expressed in around 80% of horse’s antibodies, and between 50% and 56% use IGHJ6 indicating limited use of combinations of gene segments. However, most horse antibody IgM and IgG clones (∼91%) present N-nucleotide addition, reaching 78 nucleotides in N1 and 62 in N2 regions for IgM and 111 nucleotides in N1 and 104 in N2 for IgG. These results suggest a major role of junctional diversity in generating equine antibody repertoire variability.

## MATERIALS AND METHODS

### Horse blood samples

The peripheral blood samples from four healthy, mixed male breed adult horses, aged 5 to 9 years old, were obtained in partnership with the Immunobiological Research and Production Center (CPPI) of the State of Paraná.

About 35 ml of peripheral blood was obtained from each animal using Vacutainer tubes with EDTA anticoagulant. The PBMC were isolated by Ficoll-PaqueTM gradient centrifugation. The cells (1×10^7^ cells) were cryopreserved in FBS 90%/ DMSO 10% at −80 °C until use.

The Ethics Committee approved the experimental design on the Use of Animals of the Federal University of Minas Gerais (CEUA - UFMG) under protocol number 190/2018.

### Amplification of the horse antibody BCR repertoire

Mononuclear cells (PBMCs) were isolated for RNA extraction and subsequent cDNA synthesis. Total RNA extraction was performed by the TRIzol method (Rio et al., 2010), and the RNA concentrations were verified by the Qubit RNA BR Assay kit (Thermo Fisher Scientific). According to the manufacturer’s instructions, approximately 500 ng of RNA was used for cDNA synthesis using the SuperScript IV enzyme (Thermo Fisher Scientific). The IGH amplification of the gene segments V and the constant region was carried out by multiplex PCR. A set of forward specific primers (F) for the heavy chain variable region (Manso et al., 2019) was used with new reverse specific primers (R) for the heavy chain constant region designed in this study: IgM isotype 5’ ATGACGTTGGGTAGGAAGTCCCG 3’ and IgG isotype 5’ CCACCGTGGMGTCAGAYGTG 3’. All primers have incorporated the Illumina overhang adaptors sequence to prepare the Illumina library.

Multiplex PCR reactions were conducted to obtain IGH amplicons from each of the four horses. All reactions were prepared to contain 10X High Fidelity buffer, 50 mM MgSO4, 10 mM dNTPs, 0.5 µM of each F primer, 0.5 µM of each R primer, and 0.5 U Taq DNA polymerase Platinum High Fidelity (Thermo Fisher Scientific). The cycling parameters were 94 °C for 2 min; 4 cycles of 94 °C for 1 min, 50 °C for 1 min and 72 °C for 1 min; 4 cycles of 94 °C for 1 min, 55 °C for 1 min and 72 °C for 1 min; 26 cycles of 94 °C for 1 min, 63 °C for 1 min and 72 °C for 1 min, and 72 °C for 7 min. The amplifications were analyzed on 1% agarose gels and stained with Sybr Safe (Invitrogen). The bands were excised, and purified with PCR clean-up Gel extraction (NucleoSpin).

### Library preparation and sequencing

The purified cDNAs were quantified by Qubit DNA High Sensitivity kit (Thermo Fisher Scientific). Then, it was used for sequencing libraries prepared by the Nextera XT DNA Library Prep kit (Illumina) according to the manufacturer’s instructions. The P5 and P7 indexes and adapters were incorporated into the 500 bp amplicons by the overhang adapters added to the primers. The library concentration was verified using Qubit DNA High Sensitivity kit (Thermo Fisher Scientific), and the size and quality of amplicons were confirmed with the Bioanalyzer High Sensitivity DNA Analysis (Agilent).

The IGH samples (18 pM) from the four equines were sequenced using Illumina MiSeq platform 2 × 300 bp read length.

### Bioinformatic analysis of the horse immunoglobulin heavy chain (IgH) variable-region repertoire

The reads were preprocessed by the pRESTO pipeline (vander Heiden et al., 2014), and the IG genes were annotated using IMGT/HighV-QUEST (Alamyar et al., 2012). The unproductive V(D)J rearrangements were eliminated from the dataset, as well as the productive sequences containing insertions, deletions (indels), or stop codons in V- and J-gene segments. The sequences with the same VJ segment and identical CDR H3 size were grouped using the IMGT/StatClonotype (Aouinti et al., 2016) for clonotype analyses.

After processing the sequences, analyses of the antibodies diversity were conducted, evaluating the frequencies of gene segment usage, gene subgroups, and the combination of V(D)J genes in each animal using IMGT/StatClonotype. The size, composition, and amino acids groups (defined by Crooks et al., 2004) of CDR-H3 amino acid sequences (numbered according to IMGT) (Lefranc et al., 2003) were analyzed using R studio. R studio was also used to get public repertoire antibodies defined as different horses containing antibodies with the same V and J and CDR3. To determine the reading frame (RF) of IGHD genes, we first determined the hydrophobicity index, according to Kyte-Doolittle scale, of each frame using R studio peptide package. The most hydrophilic reading frame was definied as RF1, the most hydrophobic as RF2, and the one hydrophobic with stop codons was defined as RF3 (Ivanov I, Link J., Ippolito G.C., n.d.)(ref definicao).

Other parameters such as somatic hypermutation and junction were analyzed from data extracted from IMGT/HighV-QUEST.

### Rarefaction analysis and constructing species-richness curves clonotypes

We used the program iNEXT (Hsieh et al., 2016) to subsample populations of clonotypes from immunoglobulin heavy chains that belonged to four horses based on the frequency of their occurrence in productive reads. iNEXT was also used to extrapolate beyond the number of experimentally observed productive reads that we might expect with additional sequencing. Recon (Kaplinsky & Arnaout, 2016)was used to estimate the number of missing clonotypes in the immunoglobulin heavy-chain datasets

### Statistical analysis

To compare antibody isotypes (IgG and IgM) differences, we used the Shapiro-Wilk test followed by the Mann-Whitney test.

For all analyses, the media of each horse-specific parameter followed by the average media of all four horses, and also the standard deviation of the average media of the 4 horses were used.

## RESULTS

### Restricted VJ gene usage in horse antibodies

We analyzed IgG and IgM variable heavy chains from four individual horses. Overall, the mean of raw reads per IgM samples was 1,082,148 (354,598-1,558,035) and 347,302 (210,964-481,150) for IgG samples. We obtained 31-64% of productive reads and between 40,018 to 328,300 horse antibody clones (Table 1).

**Table 1:** Overview of the IgM and IgG heavy chain variable sequencing results from four non-mmunized horses.

| **Horse**<br>**Sample** | **Ig**<br>**Isotype** | **Raw**<br>**Reads** | **Preprocessed**<br>**Reads** | **Annotated**<br>**Reads** | **Productive**<br>**Reads** | **Clones** |
| --- | --- | --- | --- | --- | --- | --- |
| 1 | IgM | 1,558,035 | 779,236 | 526,221 | 484,236 | 294,600 |
| 2 | IgM | 1,499,997 | 900,306 | 891,250 | 790,308 | 328,300 |
| 3 | IgM | 915,963 | 690,915 | 529,584 | 485,888 | 283,600 |
| 4 | IgM | 354,598 | 268,020 | 203,623 | 193,644 | 86,177 |
| 1 | IgG | 210,964 | 187,229 | 182,918 | 175,686 | 43,861 |
| 2 | IgG | 288,611 | 233,147 | 219,402 | 209,248 | 40,018 |
| 3 | IgG | 408,483 | 306,844 | 304,394 | 271,204 | 91,136 |
| 4 | IgG | 481,141 | 390,341 | 382,438 | 361,320 | 89,551 |

The EquCab3.0 horse’s genome includes 21 IGHV, 16 IGHD, and 6 IGHJ functional gene segments, leading to 2,016 possible germline coding antibodies. However, in our study, we observed a strong preference for IGHV4 subgroup gene segments, where the IGHV4-21, IGHV4-29, and IGHV4-22 are used by 80% of the horse antibodies in both IgM and IgG isotypes (Figure 1A). In addition, only 13 (of which three are present in less than 0.1% of the antibodies) from 21 IGHV seem to be used in horse antibodies.

**Figure 1:**
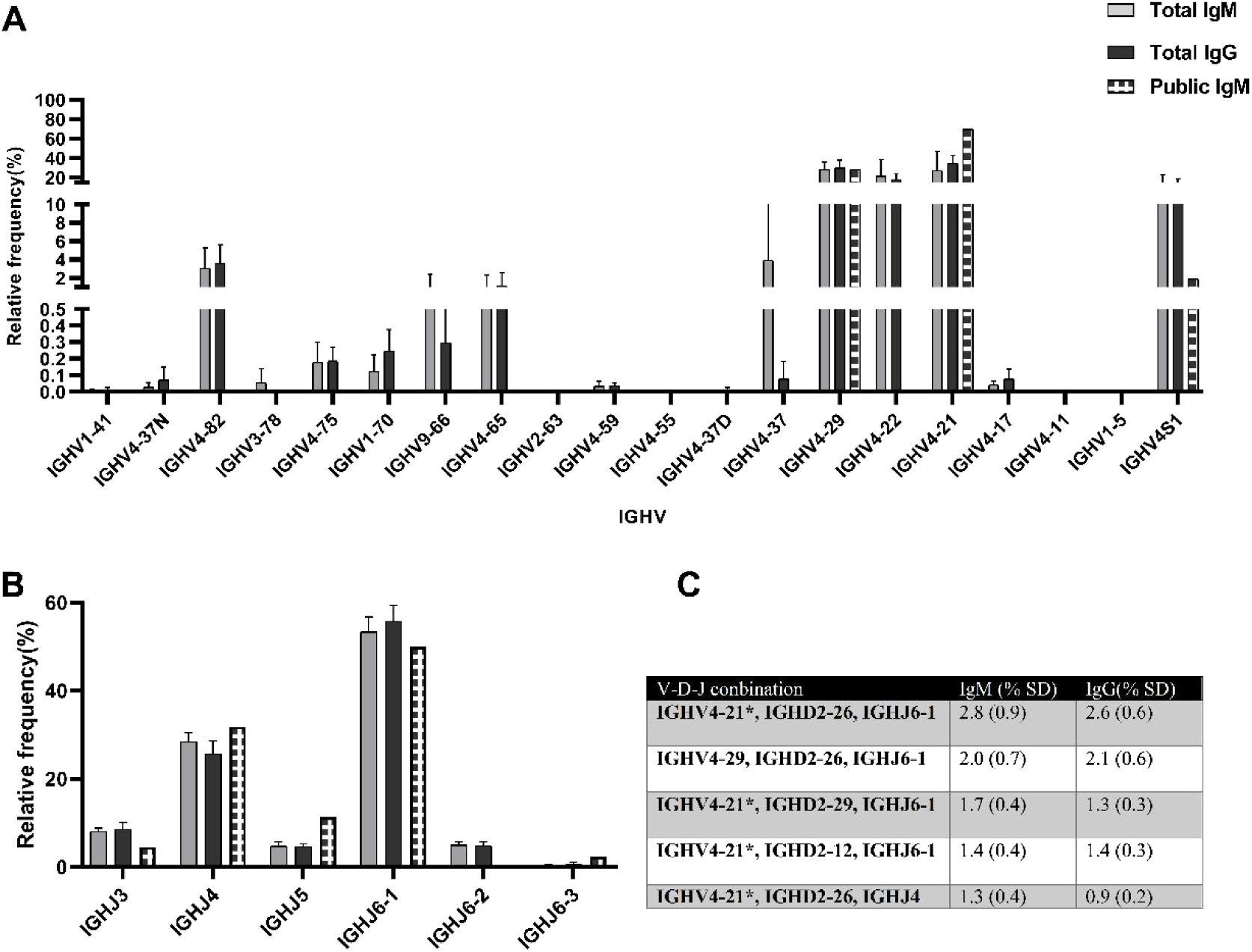
IGHV and IGHJ gene segments frequency present in public IgM and total IgM and IgG horses’ antibodies. Median of relative frequency (%) of IGHV (A), IGHJ (B) and V(D)J more frequent combination (C) in IgM and IgG isotype from four horses. The genes are organized in the order that it appears in the EquCab 3.0 genome (5’-3’).

Similarly, IGHJ4 and IGHJ6 are the preferred J gene used by both IgM and IgG isotypes (Figure 1B). Interestingly, IGHJ6 is present in almost 60% of all horse antibody clones, showing a restricted use of IGHV and IGHJ gene segments. All the 16 functional IGHD genes seem to be used by horses’ antibodies (Supplementary Figure 1).

The most frequent VDJ combinations used by horse antibodies were IGHV4-21, IGHD2-26, and IGHJ6-1, found in 2.8% (±0.9) of the IgM and 2.6% (±0.6) IgG isotypes (Figure 1C).

Our analysis also showed that more rare clones were observed in IgM samples than in IgG since the rarefaction curves do not begin to plateau, indicating that we were unlikely to capture this population’s full diversity (Figures 2A and 2B). However, we were able to capture between 52 to 66% and between 62 to 82% of all IgM and IgG, respectively (Figure 2C), with no statistical difference between IgM and IgG horse antibodies diversity compared to Shannon’s (Figure 2D) and Simpsons test (Figure 2E).

**Figure 2:**
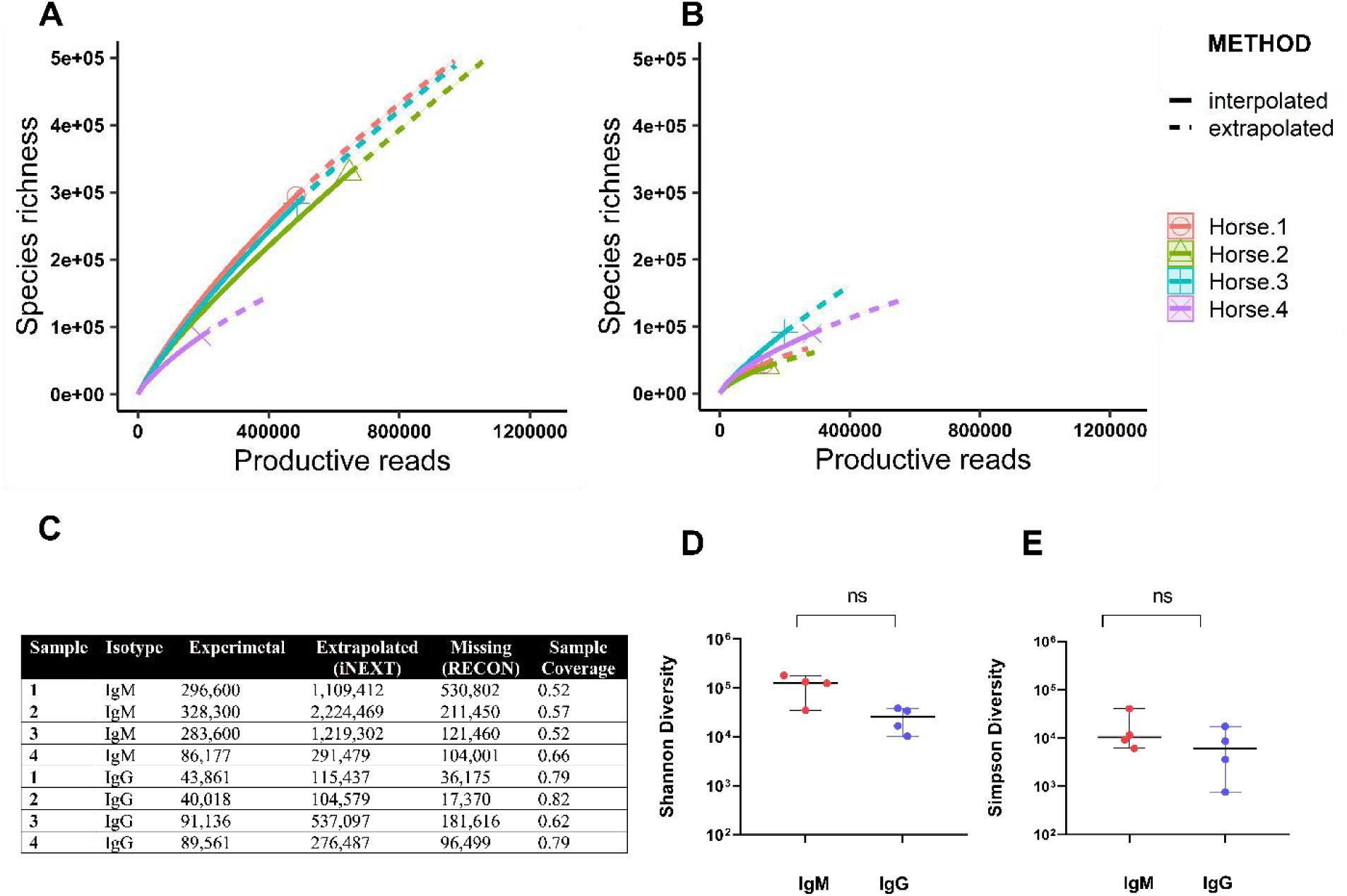
Antibody heavy chain repertoire richness and diversity estimates for IgM and IgG in the four non-immunized horses. Interpolation and extrapolation of species richness were obtained using iNEXT for IgM (A) and IgG (B). Solid lines correspond to the interpolation (based on experimental data), and the dashed lines belong to the extrapolated data. Summary of estimates for repertoire size, including missing clones (C). The comparison of Shannon’s (D) and Simpsons (E) diversity between the IgM and IgG isotypes (p < 0.05).

### Public horse antibody repertoire is enriched in shorter CDR-H3

In this study, we observed that the four horses shared (public repertoire) shared only 0.05% (44 clones) of their IgM repertoire and 0.0099% of the IgG repertoire (4 clones) (Figures 3A and 3B). For the IgM public repertoire, most of the clones present the IGHV4-21 gene (77%) of the IGHV genes in the IgM public repertoire, while in the total repertoire, it represents approximately 32% (Figure 1A). In the case of IGHJ genes, we observed an increased gene usage of IGHJ4 (from 4% to 9%) and IGHJ5 (from 29 to 32%) in the public IgM repertoire (Figure 1B).

**Figure 3:**
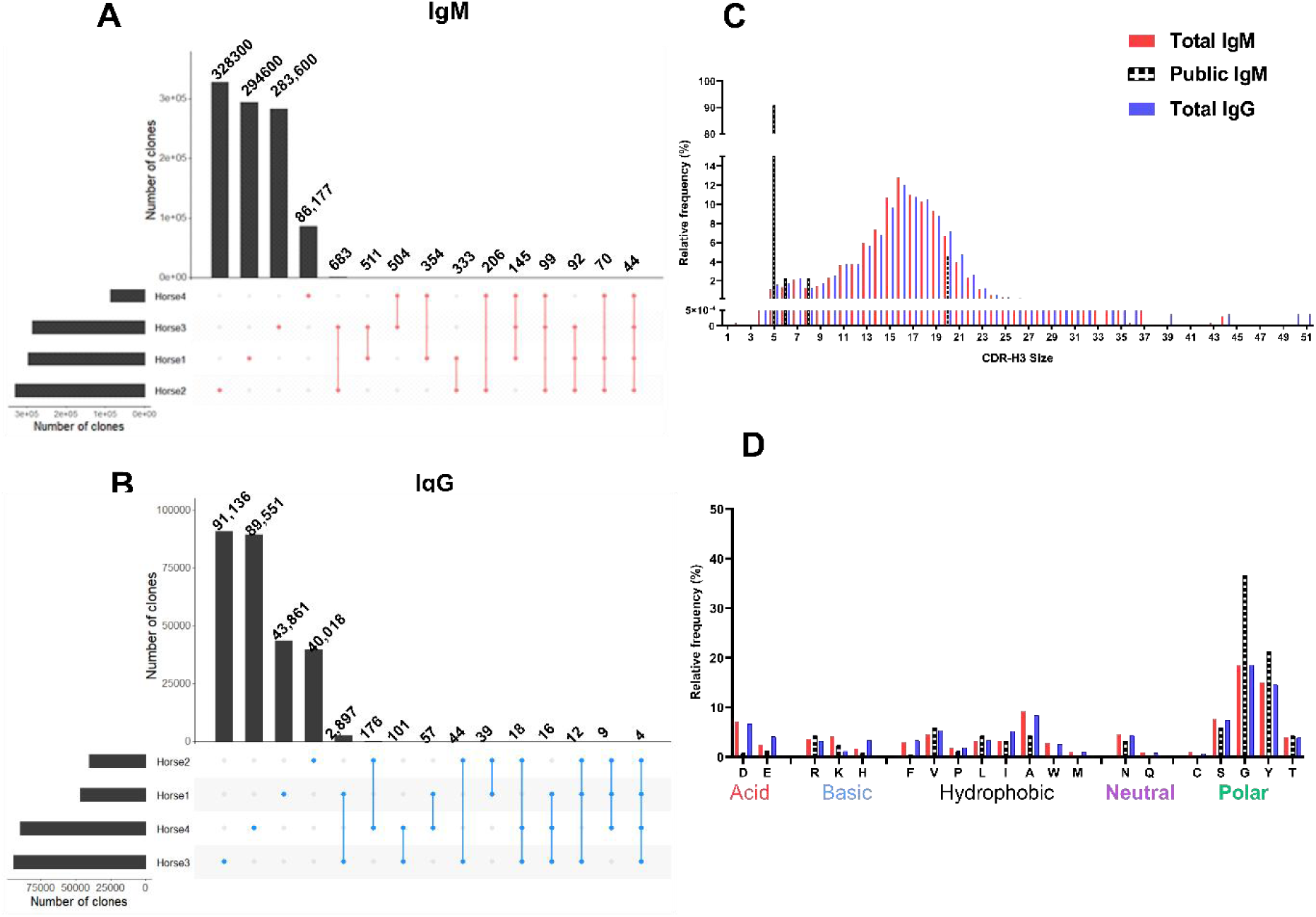
Horse public and private heavy chain variable region repertoire. The number of antibody clones presented by the different horses and the shared number of clones between the 2, 3, or 4 horses in this study for the IgM (A) and IgG (B). Comparison of the CDR-H3 length (C) and amino acid composition (D) between the Total IgM, Total IgG, and the public IgM repertoire.

Interesting to note that more than 90% of the CDR-H3 found in the IgM public antibody repertoire presented only five amino acids length (Figure 3C). In general, the CDR-H3 size distribution of horses follows a bi-modal pattern, with sizes ranging from 4 to 51 amino acids residues with a median length of 16 residues for both IgM and IgG isotypes (Figure 3C).

Interestingly, polar amino acids such as glycine (G) and tyrosine (Y) are increased in IgM public repertoire, differently from the acidic aspartic (D) and glutamic acids (E) and the hydrophobic phenylalanine (F), tryptophan (W) and alanine (A) that decreases (Figure 3D).

Our results suggest that the horse IGHV repertoire appears to be derived from limited germline gene families.

### Characterization of somatic hypermutation (SHM) frequency and pattern found in horse antibodies

Based on our previous results, we hypothesized that the biggest horse immunoglobulin diversity comes from somatic hypermutation and junctional diversity.

Here, we observed that the frequency of mutations in IgG isotype (media: 7.22%) was similar to IgM (media: 6.46%) compared to their germline mapped on EquCab3.0 genome (Figure 4A). The majority of mutations were found in CDR regions, especially at positions 32 (CDR1), 50, 52 and 58, from CDR2 and 88 (FR3) for both IgM and IgG isotypes (Figure 4B). We also observed that an average of 16.79% of nucleotides are mutated in CDRs of region IGHV, from which the majority of them (45 to 51%) are present in AID motifs (RGYW and complementary WRCY nucleotide motifs) (Supplementary Table 1).

**Figure 4:**
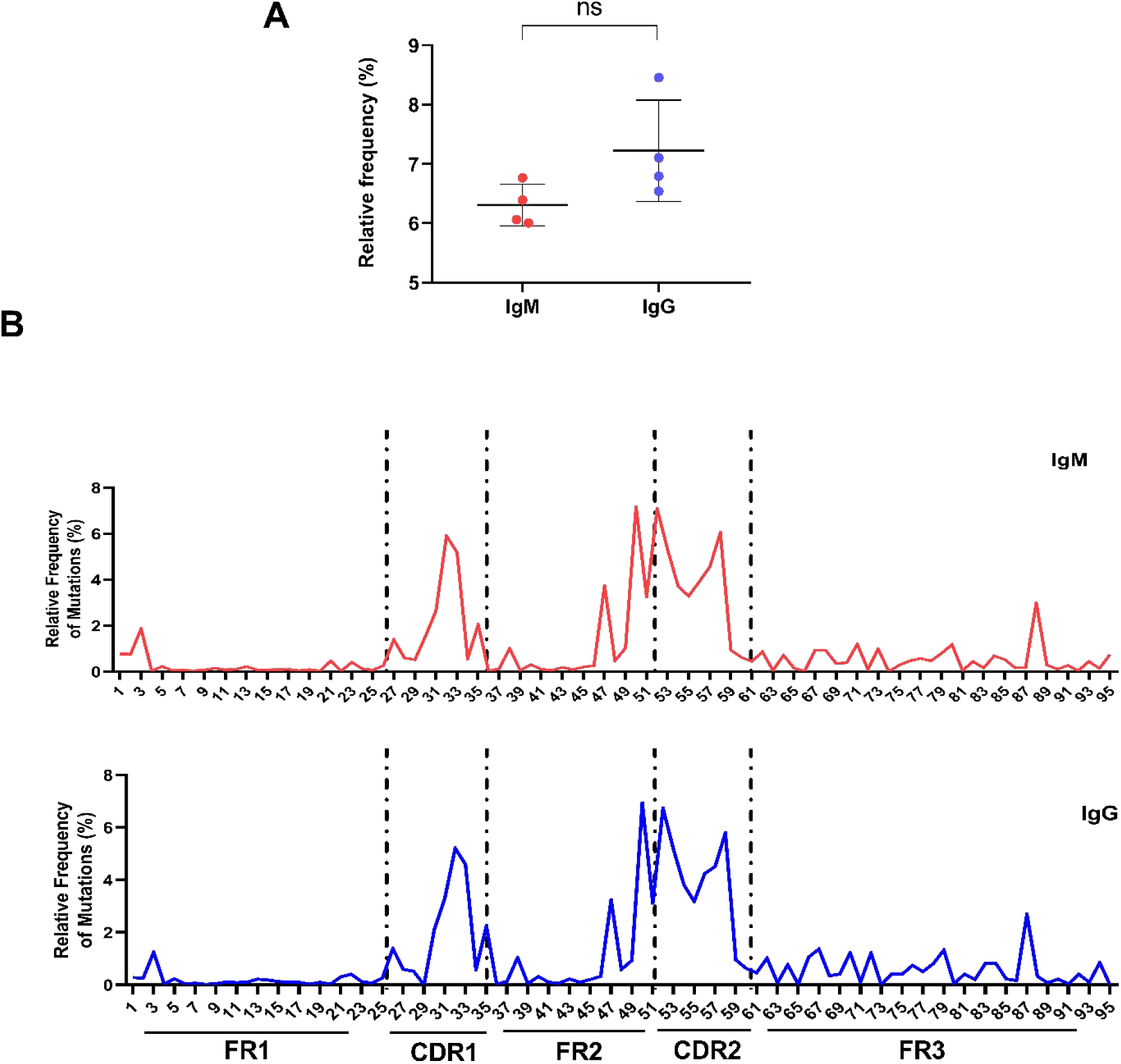
Somatic Hypermutation (SHM) characteristics of Horse IgG and IgM heavy chain variable region. (A) Media of SHM frequency (%) at the IGHV gene segment from IgM and IgG isotypes (p < 0.05). (B) The number of mutations by amino acid position in the IGHV gene segment of the horses’ heavy chain (According to the IMGT numbering, without gaps). The dotted lines delimit the FR and CDR regions. IgM is shown in red and IgG in blue.

### Characterization of Horse Antibodies Junctional Diversity

An essential source of antibody diversification is the addition and deletion of nucleotides between VDJ junctions. Therefore, we analyzed the occurrence of the P/N nucleotide addition and exonuclease trimming for both IgM and IgG horse antibodies. We observed very similar characteristics in all the junctional regions of IgM and IgG horse antibodies (Figures 5A and 5B). N1 and N2 nucleotides media vary from 8.6 to 9.2 present in around 92 % of the IgM and IgG antibody clone (n = 998,756 clones for IgM, n = 264,566 clones for IgG (Figura 5A and 5B, Table 2). Interestingly, 43-44% of the antibodies have N1 (ranging from 10 to 111 nt) and N2 junctions (ranging from 10 to 104 nt) with 10 or more nucleotides, and 6.9 to 9.8% of these regions with 22 or more nucleotides (Table 2). It was also possible to observe that half of the 10 biggest CDR3 present cysteines (Supplementary table 4). We also noticed that N1 region is highly enriched in G (35.59%), and the N2 region is enriched in G and T (30%) for both isotypes (Supplementary Table 2).

**Figure 5:**
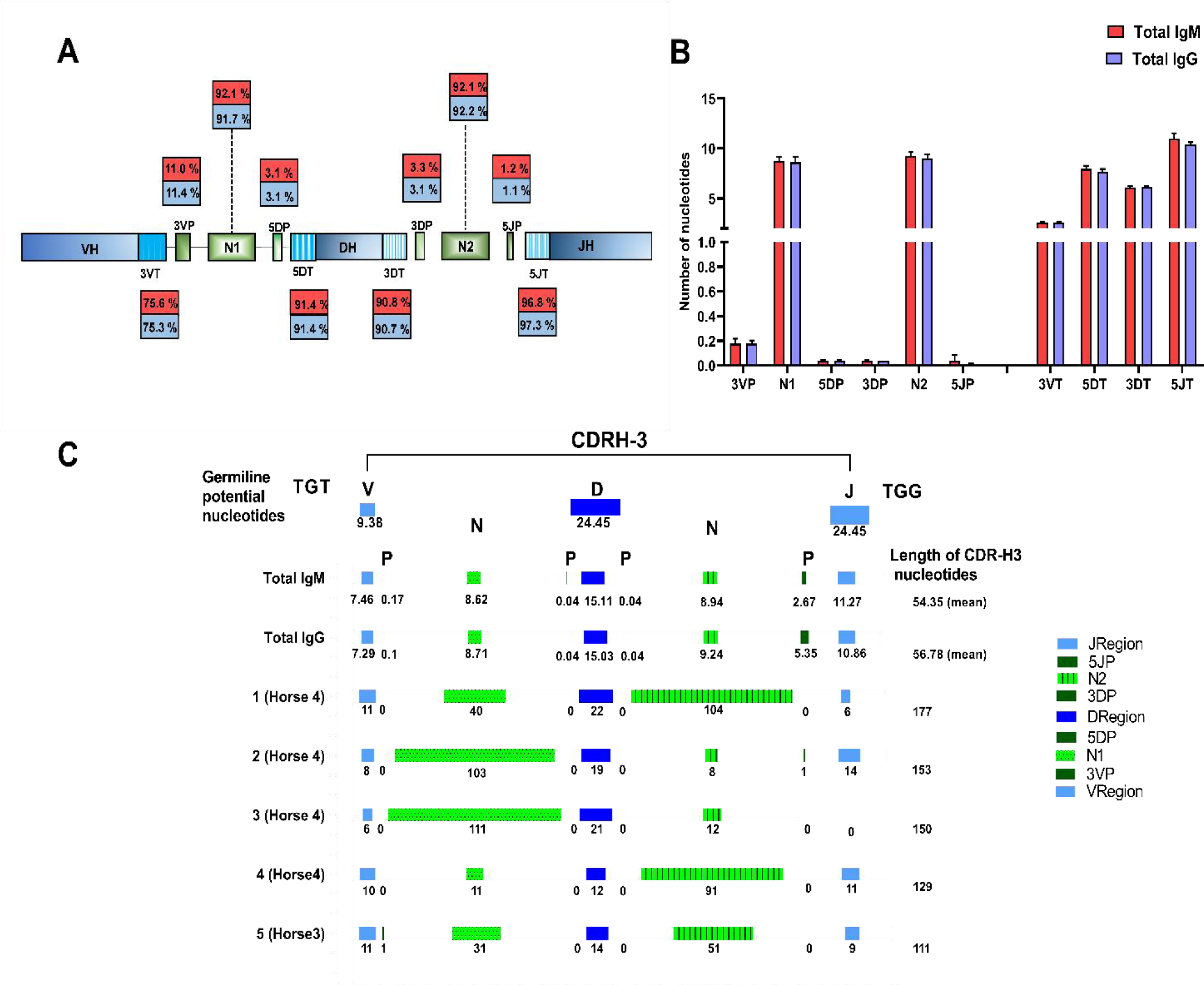
VDJ junction analysis. (A) Junctional modifications schema during VDJ rearrangement, showing the locations and the occurrence of different types of junctional modifications. Into the box are the mutation frequency (%) of each junctional segment in all the antibodies clones analyzed: 3VP and 3VT for 3′V region, 5DP and 5DT for 5′D genes, 3DP and 3DT for 3′D region, and 5JP and 5JT for 5′J genes, where P means palindromic nucleotides additions, and T means exonuclease trimmings; N1 the non-template randomized nucleotides additions at the 3′V and the 5′D genes; N2 for N additions at the 3′D and the 5′J genes. (B) The median number of nucleotides per junction region added or trimmed.(C) Deconstruction of the components that contribute to the length of the CDR-H3 in the media of total of the IgM and IgG clones, as well as the 5 biggest CDR-H3 of the IgG clones. The mean of nucleotides of the germline sequence of the VH gene segment, P and N junctions, the DH gene segment, and the JH gene segment to the CDR-H3 length is illustrated.

**Table 2:** Analysis of nucleotide additions in Horse Antibodies.

| Sample | Number of clones | Mean number of N1 nucleotides (range) | Mean number of N2 nucleotides (range) | Clones with N1 additions longer than 10 nucleotides/ and longer than 22 nt (%) | Clones with N2 additions longer than 10 nucleotides/and longer than 22 nt (%) |
| --- | --- | --- | --- | --- | --- |
| IgM | 998,756 | 9.54 (0-78) | 9.71 (0-62) | 43.95/7.4 | 43.74/8.6 |
| IgG | 264,566 | 8.72(0-111) | 9.24 (0-104) | 43.67/7.6 | 45.17/10.0 |

**Table 3:** Amino acid percentage composition per reading frame of horse D gene.

|  | **RF1** | **RF2** | **RF3** | **iRF1** | **iRF2** | **iRF3** |  |
| --- | --- | --- | --- | --- | --- | --- | --- |
| **D** | **5.22** | 0.75 | 0 | 0 | 0 | 0 | **Acid** |
| **E** | 0.74 | 0 | 2.90 | 0 | 0 | 3.00 |  |
| **R** | 0.74 | 1.50 | 4.34 | 5.97 | 0.74 | 0 | **Basic** |
| **K** | 0.74 | 0 | 0 | 0.74 | 0 | 0 |  |
| **H** | 0 | 0 | 0.72 | 21.64 | 1.49 | 0 |  |
| **F** | 0 | 0 | 0.72 | 0 | 0 | 0.75 | **Hydrophobic** |
| **P** | 0 | 2.25 | 0 | 2.24 | 4.48 | 17.29 |  |
| **V** | 0 | 27.82 | 0.72 | 0.74 | 29.10 | 3.00 |  |
| **L** | 0 | 5.26 | 42.75 | 1.49 | 1.49 | 18.04 |  |
| **I** | 2.23 | 6.77 | 1.44 | 0.74 | 18.66 | 3.00 |  |
| **A** | 5.97 | 3.00 | 1.45 | 3.54 | 5.26 | 2.93 |  |
| **W** | 3.73 | 0 | 13.76 | 0 | 0 | 0 |  |
| **M** | 0 | 18.04 | 0 | 0 | 0 | 0.75 |  |
| **N** | 5.97 | 6.66 | 0.72 | 15.67 | 2.99 | 0 | **Neutral** |
| **Q** | 0 | 0 | 2.90 | 1.49 | 0 | 4.51 |  |
| **C** | 0.74 | 0 | 5.80 | 5.97 | 0 | 2.94 | **Polar** |
| **S** | **14.92** | 0 | 0 | 23.88 | 4.48 | 6.77 |  |
| **G** | **18.65** | 4.5 | 1.45 | 0 | 5.97 | 1.50 |  |
| **Y** | **39.55** | 0 | 2.90 | 11.94 | 0.74 | 2.25 |  |
| **T** | 0.74 | 29.32 | 0 | 0.74 | 26.12 | 1.50 |  |
| **\*** | 0 | 0 | **17.39** | 4.47 | 0.74 | **32.33** |  |
| AA Media by RF | 7.88 | 7.82 | 8.11 | 7.88 | 7.88 | 7.82 |  |

Similarly, exonuclease trimming was observed in around 70% to 97% of the Ig clones, with the biggest number of nucleotides trimmed in the IGHJ gene ends (mean of 10 for IgM and 11 for IgG) (Figure 5A and 5B). When analyzing the components that contribute to the length of the CDR-H3, we observed that an average of 15 nucleotides from IGHD gene segment contribute to the length of the IgM and IgG CDR3s with contain an average of 54 nucleotides (around 26-27% of IGHD contribution to the CDR3). Surprisingly, when the 5 biggest CDR-H3 of the IgG clones where anlysed we observed a contribution of 12 to 22 nucleotides of IGHD genes which represents only 9% to 12% of the CDR3. The biggest contribution in these cases came from N1 addition that can contribute with 111 nucleotides of a CDR3 with 150 nucleotides (74%) or the N2 addition that can contribute with 91 nucleotides of a CDR3 containing 129 nucleotides (70%) (Figure 5C).

We also observed a preference for the use of RF1 (more of 80%) in the horse’s antibodies (Supplementary Figure 2), enriched in polar amino acids such as tyrosine and glycine (Table 2). Of the 96 possible sets of IGHD amino acid sequences (16 x 6 = 96), 36 (37.5%) include one or more tyrosine, while only 13 (13.5%) have one or two arginines (Supplementary Table 3).

## Discussion

In this work, we investigate the antibody-heavy chain repertoire of four different horses, presenting the largest collection of adaptive immune receptor sequences described to date for horses. We analyzed 40,018 to 328,300 horse IgG or IgM clones, with good coverage, from 52 to 82% of the repertoire. Similar to this work, it was observed a difference in depths for IgM (36%) when compared to IgG (64%) for human antibodies (Galson et al., 2015). Although not much difference was observed between IgM and IgG isotypes, this is the first high-throughput sequencing study that characterizes both isotype’s horse repertoires.

Interestingly, we found that approximately 80% of the IgM and IgG antibodies present the IGHV4 group as a gene segment used in their antibodies, corroborating works from, Similar results were also observed by Tallmadge and collaborators (2013) using 5’ RACE, in which they found a strong preference (80%) for the IGHV4-29 (previously called IGHV2S3) and IGHJ6-1 (55%) (previously called IGHJ1S5) usage in adult horses’ antibodies (Chaudhary & Wesemann, 2018; Manso et al., 2019; Sun et al., 2010; Tallmadge et al., 2013). Similar, humans antibodies also have a preference for the IGHV4 family (Arnaout et al., 2011), differing from other organisms like cattle, dogs, and mice that present a predominance for IGHV1 genes in their antibodies and cats with the presence of IGHV3 genes(Pasman et al., 2017; Rettig et al., 2018; Steiniger et al., 2014, 2017). We also observed a predominance of IGHJ6 in horse antibodies, while dogs and cats antibodies se mostly IGHJ4, and mouse, the IGHJ1 group (Arnaout et al., 2011; Steiniger et al., 2014, 2017). It is interesting to note that horse antibody repertoire is highly dominated by only a few IGHV and IGHJ genes, even if they can use all of their theoretical germline combinations. We observed a high frequency of antibodies (2.8 ±0.9 %) containing IGHV4-21, IGHD2-26, and IGHJ6-1 gene segments combination, also identified in previous work on non-immunized horses (Manso et al., 2019; Tallmadge et al., 2013). This result is not different from human antibody repertoires (Arnaout et al., 2011), where 0.1% to 2.7% of sequences have the same V(D)J combinations.

In addition, even for a few clones, we observed the presence of a public horse antibody repertoire in the absence of any specific immune stimulation. We found more clones in the public IgM (0.05%) repertoire than in the public IgG repertoire (0.009%). The small number of shared public antibodies clones can be due to the high diversity of horses antibodies but can also be due to an artifact of the clonotyping method used in this work that considers the same clone only antibodies with identical CDH3 region (IMGT/HighVQuest). The observation of more public IgM (1.4%) than IgG (0.3%) and IgA (0.5%) was also observed in the human antibody (Galson et al., 2015. Interestingly, even with a smaller number of public antibodies, we found differences between the CDR-H3 length of the public and the entire repertoire, observing a higher percentage of short CDR-H3 in the public repertoire, also observed in human (Briney et al., 2019; Galson et al., 2014; Soto et al., 2019). It is supposed that B cells expressing receptors with short CDR-H3 are selected because they increase their affinity for the antigen, make clonal expansion, and differentiate in plasma cells or memory B cells (Rosner et al., 2001).

In this work, we also evaluated the SHMs in the IGHV gene segment of the animals (FR1, CDR1, FR2, CDR2, and FR3 regions). The IgG sequences showed a similar mutation frequency than the IgM sequences, probably due to limited pathogen stimulation since they are not immunized. To better understand how mutations are distributed along the IGHV gene segment, we evaluated the number of mutations present in each position for both studied Ig isotypes. We observed a similar mutation profile between the IgG and IgM, with more mutations found in the CDR regions, even in these non-immunized animals. This corroborates previous studies in adult horses (Tallmadge et al., 2013) and healthy and HIV patients (Bowers et al., 2014).

The mechanism of variability that produces SHM is carried out by the enzyme cytidine deaminase (AID) induced by activation, by deamination of the cytosine base, creating a U:G mismatch. The AID targets SHM mutations on “hotspots” (complementary RGYW and WRCY nucleotide motifs) (Spencer & Dunn-Walters, 2005). This work observed that around 19% of IGHV sequences present AID motifs, similar to human IGHV, presenting 17.8% of these motifs (Bowers et al., 2014). In addition, as in the human repertoires, we found a higher percentage of motifs in the CDR (average of 13.5%) than in the FR (average of 5.2%), as well as a higher number of mutated nucleotides in this region (Bowers et al., 2014).

It is important to highlight that, besides a very similar VJ gene usage in IgM and IgG antibodies, and also a very similar profile of SHM, the IgM and IgG repertoire looks very dissimilar according to Bray-Curtis dissimilarity index (data not show).

Very little has been described the characteristics of horse junctional diversity in horse antibodies. Here, we observed N1 and N2 nucleotide additions in most IgM and IgG clones, and also observed exceptionally long N nucleotide additions for both N1 and N2 IgG (10–111 bp) in around 44% of the antibodies. Non-template additions to IGH genes have been reported in humans and mice (Shi et al., 2014), pigs (Šinkora et al., 2003), and cattle (Liljavirta et al., 2014). The mean number of nucleotide addition in humans is 6.6 ± 4.3 in N1 and 6.4 ± 4.6 in N2, while in mice, it is 2.4 ± 2.2 and 2.1 ± 1.8 in N1 and N2, respectively (Shi et al., 2014). The average number of nucleotide additions in cattle that present ultralong CDR3s is not particularly high (2.5 in N1 and 2.6 in N2), reflecting the high frequency (35%) of unions with zero additions (Liljavirta et al., 2014). Such frequency and length of N additions have not been reported in other species, suggesting that this diversity mechanism is essential to generating variability in equine immunoglobulins.

Extensive trimming of IGHD genes in horse antibodies (mean value of 6 and 7.9 nucleotides for the 3D and 5D junction, respectively) observed is not very different from cattle antibodies (trimming of 5 to 6 nucleotides) (Liljavirta et al., 2014). It is interesting to note that the larger trimming followed in horse antibodies was observed in the IGHJ gene, ranging between 10 and 11 nucleotides, while in other species, such as cattle, humans, and mice, have respectively 2, 6, and 4 nucleotides trimmed in this region (Liljavirta et al., 2014, Shi et al., 2014). In our data, between 70% to 97% of antibody clones had nucleotides deleted anywhere in the junction, similar to other species (Liljavirta et al., 2014; Shi et al., 2014).

This impressive N nucleotide addition frequency and length can be due to differences in Terminal deoxynucleotidyl transferase (TdT) enzyme activity in horses compared to other species.

These enzymes are composed of mainly two regions, a catalytic core composed of finger, palm, and thumb domains at the C-terminus and a BRCA1 (breast cancer susceptibility protein) C-terminal (BRCT) domain at the N-terminus. When comparing the horse sequence of TdT isorform 1 and 2 with human, mouse, pig, and cattle TdT we observed the conservation of the catalytic aspartic acids and the substrate-specific loop 1 (Figure 6). Interestingly, the palm domain region, between the first aspartic acids and the loop1, is one of the most different regions between the TdT from other species.

**Figure 6:**
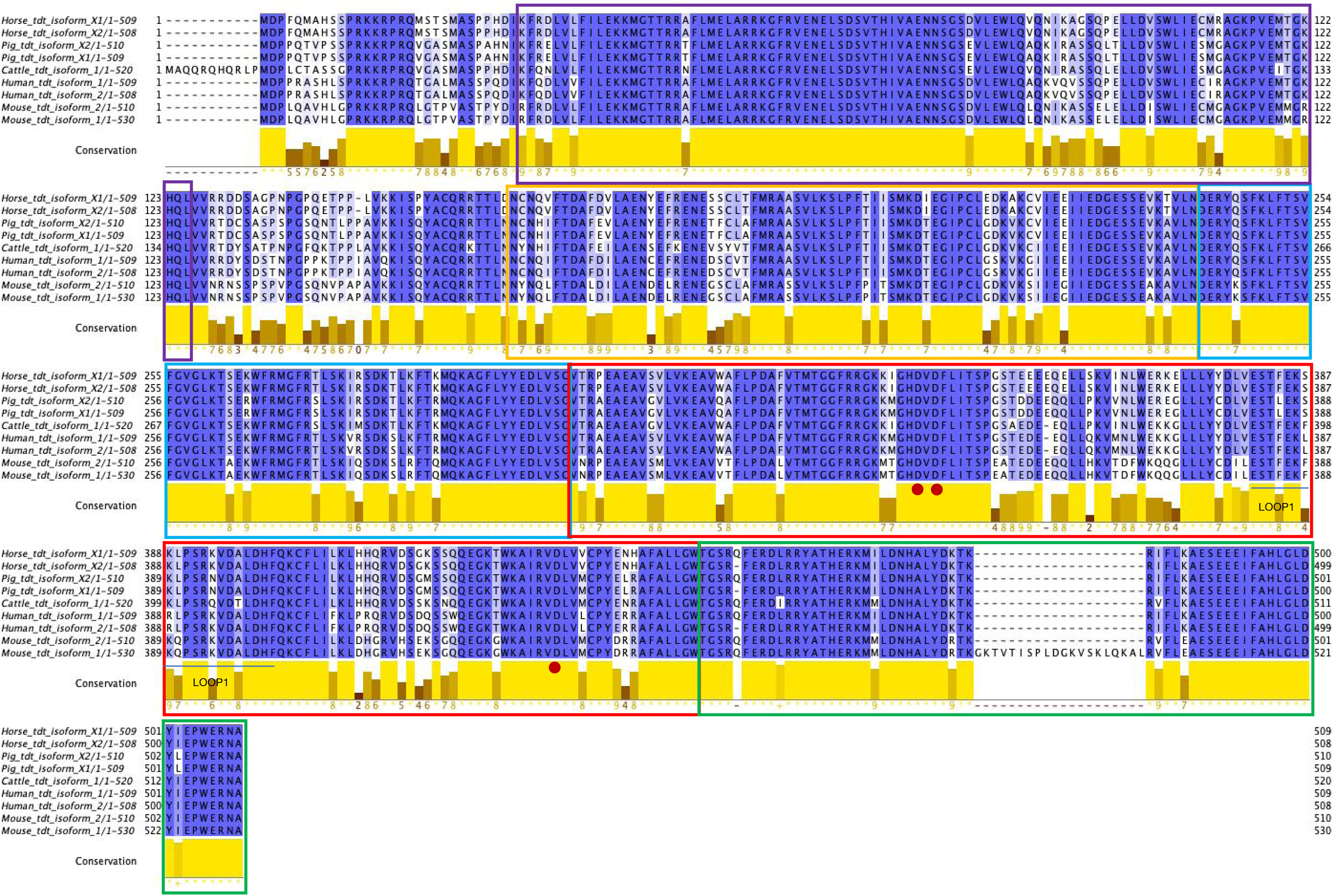
Multiple Sequence alignment of TdT from different species. An amino acid sequence alignment of the two equine TdT isoforms with other vertebrate TdT was done using the Clustal Omega program (Sievers et al., 2011) present in Jalview package (Waterhouse et al., 2009)that also provided Conservation analysis. The different box colors represent TdT domains, BRCT domain annotated by CDD (Lu et al., 2020) and TdT catalytic core as described(Delarue et al., 2002) Clique ou toque aqui para inserir o texto. : Purple: BRCA1 C Terminus (BRCT) domain (CL0459), Yellow: Helix-hairpin-helix domain (HHH_8), Blue: Fingers domain of DNA polymerase lambda, Red: DNA polymerase beta palm, highlighting the 3 catalytic Aspartic Acids (red circle) and loop1, Green: DNA polymerase beta thumb. NCBI Reference Sequences: Horse_tdt_isoform_X1 (XP_005602408.1); Horse_tdt_isoform_X2 (XP_001501812.3); Pig_tdt_isoform_X2 (XP_003133204.1); Pig_tdt_isoform_X1 (XP_005671421.1); Cattle_tdt_isoform_1 (NP_803461.1); Human_tdt_isoform_1 (NP_004079.3); Human_tdt_isoform_2 (NP_001017520.1); Mouse_tdt_isoform_2 (NP_001036693.1); Mouse_tdt_isoform_1 (NP_033371.2)

In addition to this region, we can also observe a very dissimilar region between the BRCT domain and TdT catalytic core. For the best of all knowledge, it is unclear how this non-enzymatic domain contributes to the unique biological function of TdT. Interesting to note that this interdomain region is enriched in Proline amino acids. It looks like for DNA polymerase lambda, which presents a bigger Proline-rich domain in this region compared to TdT, this region can impact DNA polymerase fidelity and with BRCT domain can act cooperatively to promote primer/template realignment between DNA strands of limited sequence homology (Fiala et al., 2006; Taggart et al., 2014). Since the TdT template and untemplated activities (Loc’h et al., 2016) are proposed to be essential for antibodies diversity, future studies need to investigate the role of the interdomain region in these activities, as well as the role of the region in palm domain in between aspartic acids and loop1.

This study also observed that more of 80% of the horse antibodies use reading frame 1 (RF1) for both the IgM and IgG isotypes. Several species, such as humans, mice, and sharks, produce antibodies using IGHD reading frame 1 (RF1) (Schroeder et al., 2010). However, similarly to other species RF1 used for horses antibodies is strongly enriched in tyrosine, representing 39.55% of the IGHD amino acids. We know that tyrosine is the amino acid that typically makes the most significant contribution to binding affinity at protein ligand-receptor interfaces (Bogan & Thorn, 1998). This suggests that natural selection was operating on immunoglobulin diversity gene segments to restrict and control their evolution in such a way as to influence the composition and range of diversity of immunoglobulin antigen-binding sites (Burnet, 1976).

## Conclusions

This is the first high-throughput sequencing study that characterizes IgM and IgG isotype horse repertoire to the best of our knowledge. We showed a highly restrict use of IGHV and IGHJ genes in horse antibody repertoire in which around 80% of the antibodies are composed by only 3 IGHV gene segments and almost 60% of them with the same IGHJ gene segment. We observed a complex and diverse repertoire for IGH, given mainly by the junctional diversity, much bigger and frequent than the one present in other species. Our study on the equine antibody repertoire contributes to understanding the generation of their diversity and open up new questions about horse TdT particularities to generate such diversity.

## Author Contributions

CN: designed the study, did all the experiments, analyzed the data and wrote the paper; TM: designed and validated primers, designed the study and discussed the results; FM: help in data analysis; LM: helped in the initial analysis of the data; RM: helped Illumina library preparison and sequenced the samples; JM, BA: collected horse samples; AV: discussed the results; JMD, GI: help primer design, study design and wrote the paper; LFF: designed the study, discussed and analyzed the data and wrote the paper

## Acknowledgments

SynBiom group for fruitful discussions, specially Dr. Marcella Nunes de Mello-Braga and Dra. Marcele Rocha Neves Rocha. A special acknowledge to Regina Maria Fernandes for project management.

---

**Supplementary Table 1:** AID (RGYW/WRCY) motifs and targeted mutation frequencies in CDR and FR regions.

|  | IgM | IgG |
| --- | --- | --- |
| **Number of RGYW/WRCY motifs per IGHV segment** | 19 (1.18) | 20 (1.17) |
| **% of CDR nucleotides mutated** | 16.79 | 18.81 |
| **% of CDR mutations present in RGYW/WRCY motifs** | 51.61 | 45.85 |
| **% of FR nucleotides mutated** | 4.24 | 4.71 |
| **% of FR mutations present in RGYW/WRCY motifs** | 16.92 | 18.76 |
| **% of all nucleotides mutated** | 6.37 | 7.12 |
| **% of all mutations present in RGYW/WRCY motifs** | 24.20 | 25.59 |

**Supplementary Table 2:** Percentage of nucleotides present at the N1 and N2 junctions of horse IgM and IgG antibodies.

|  | %A | %T | %G | %C |  |
| --- | --- | --- | --- | --- | --- |
| **IgM** | 24.82 | 20.21 | **35.59** | 19.37 | **N1.REGION** |
| **IgG** | 24.28 | 21.47 | **34.90** | 19.35 |  |
| **IgM** | 21.29 | **28.20** | **30.32** | 20.19 | **N2.REGION** |
| **IgG** | 21.57 | **29.60** | **28.87** | 19.96 |  |

**Supplementary Table 3:**
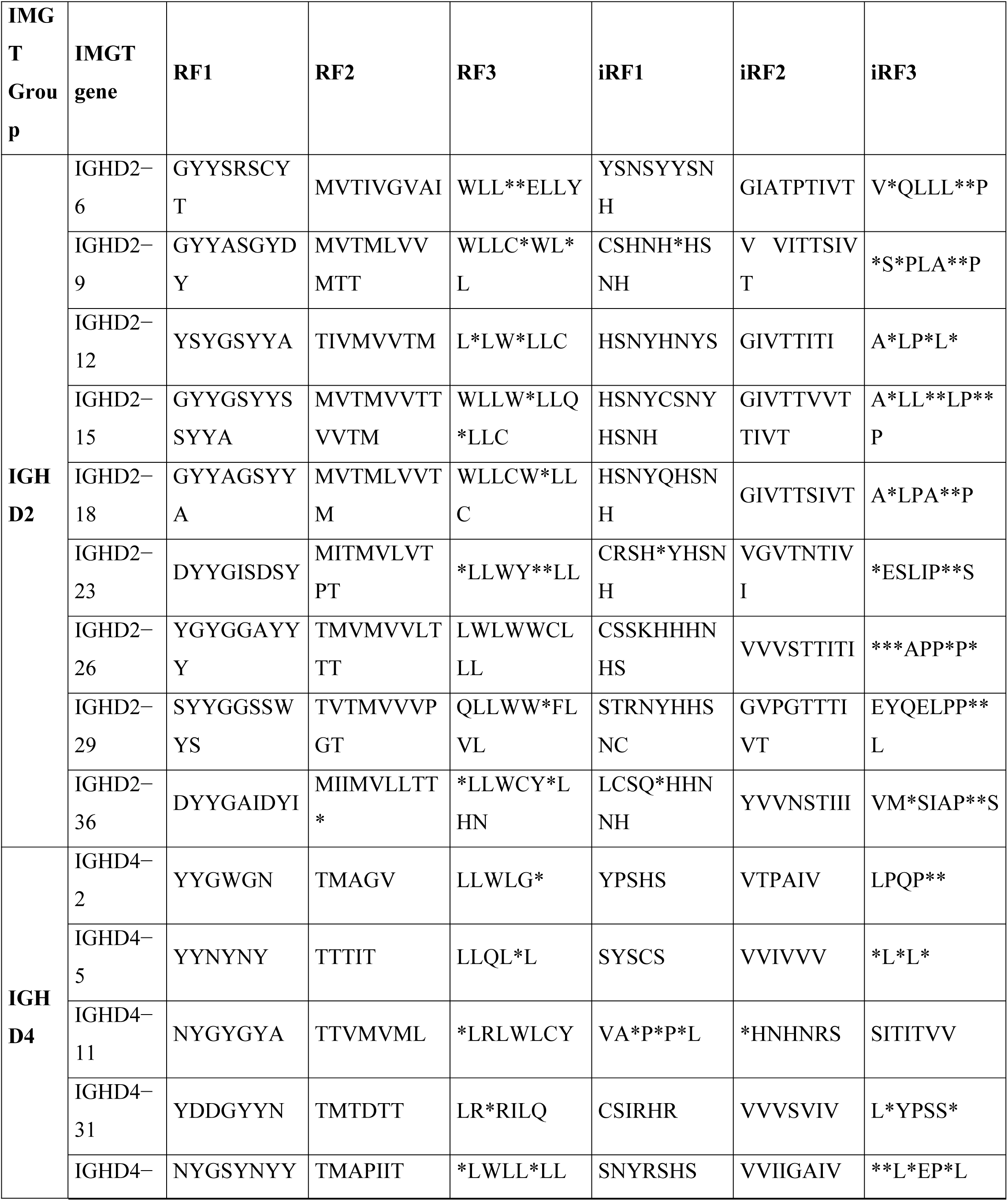
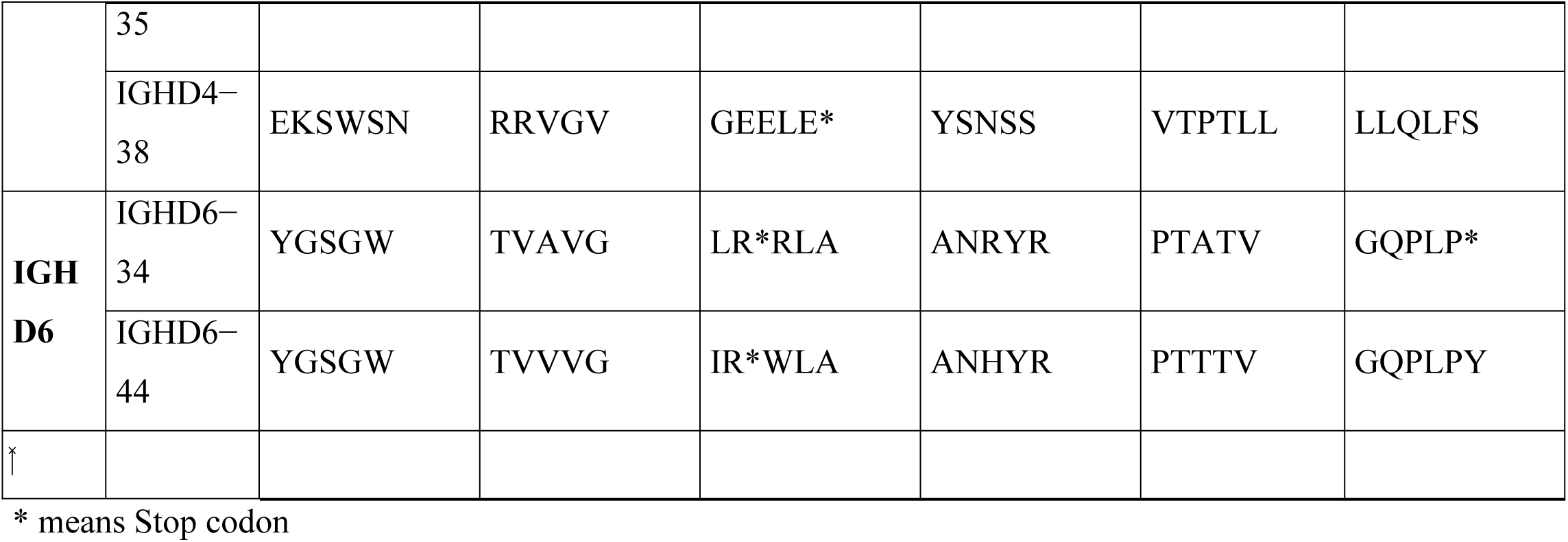
Amino Acid Composition per reading frame of IGHD functional gene segments in horse antibodies.

**Supplementary Table 4:**
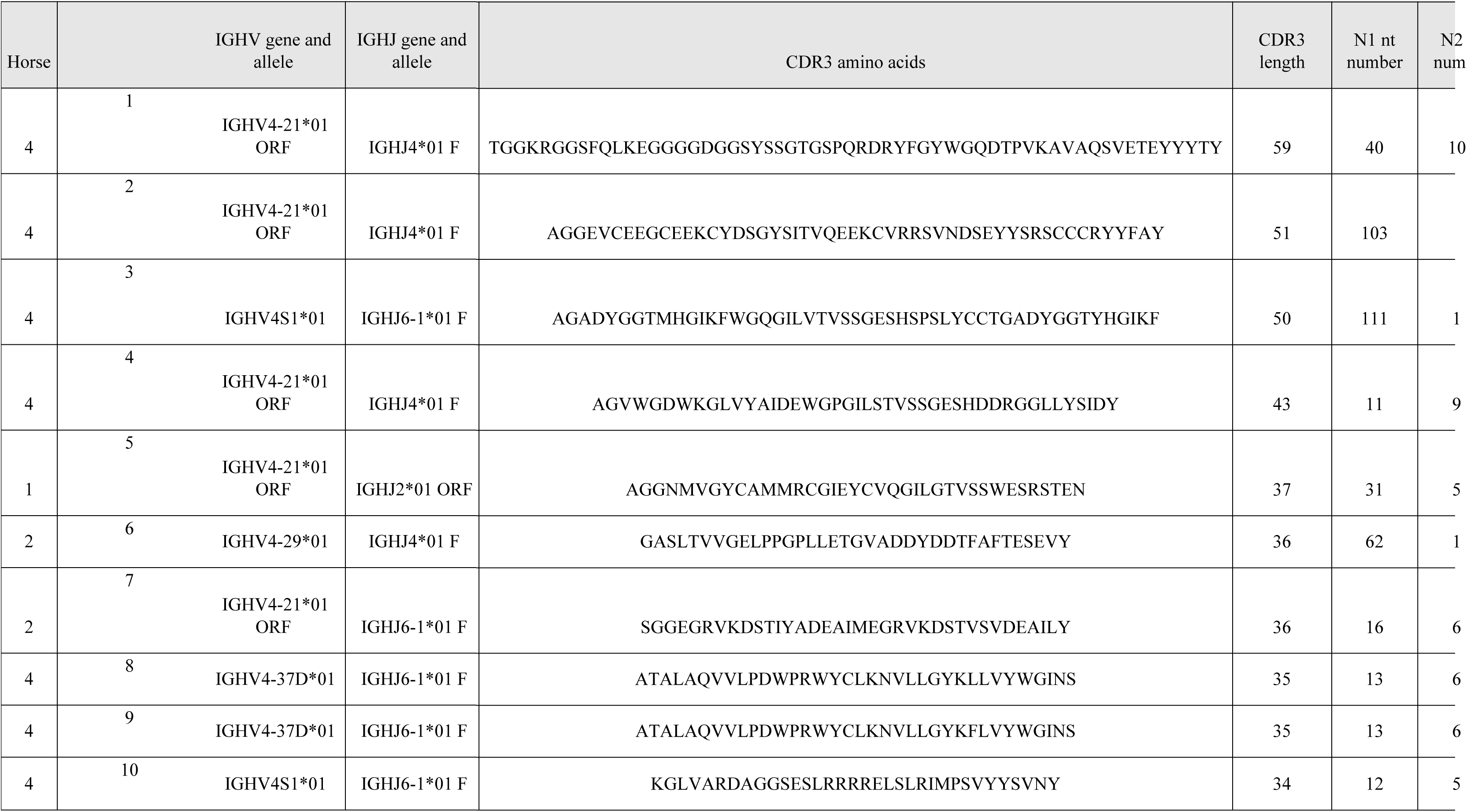
Top 10 bigger CDRH3 found in this study.

**Supplementary Figure 1:**
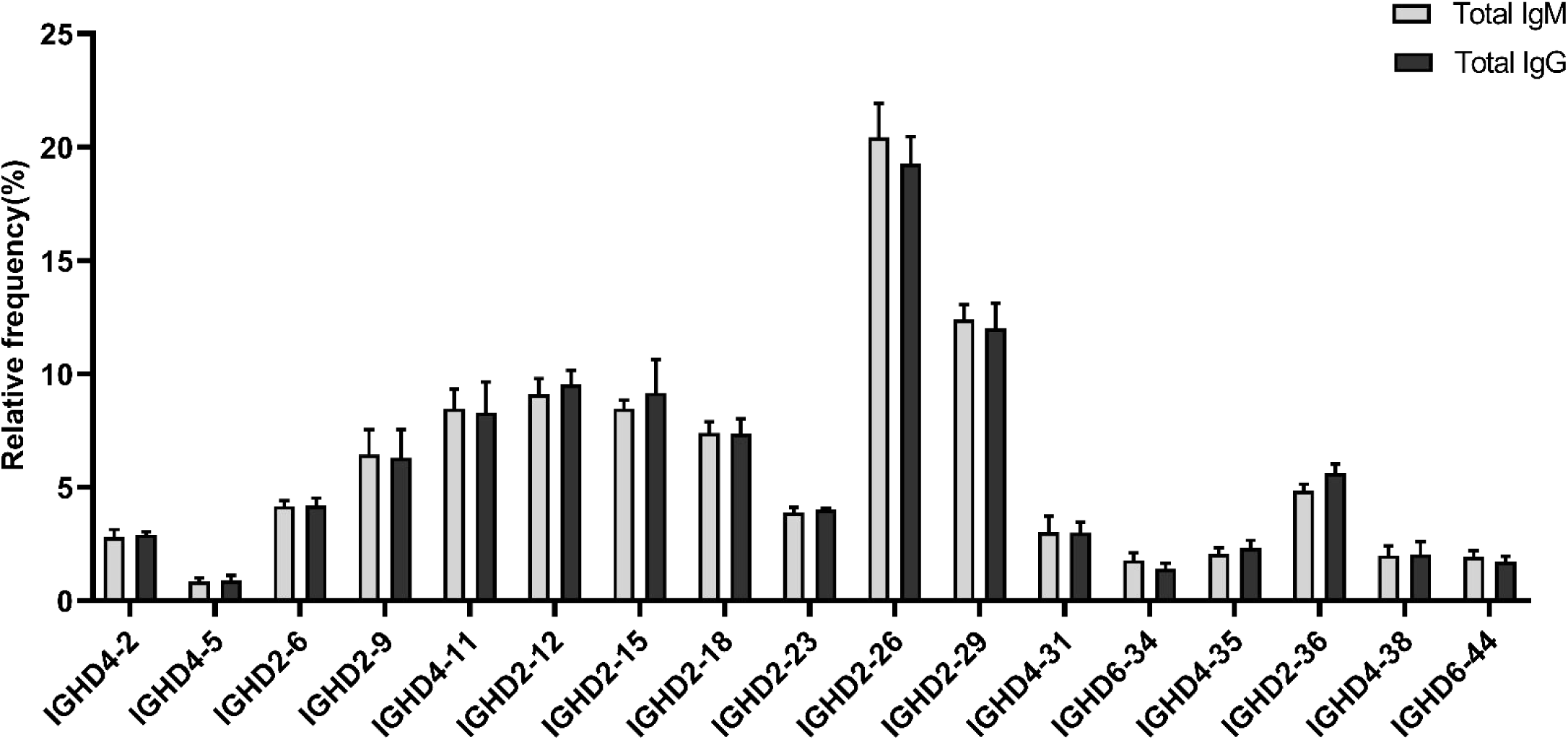
Median of Relative frequency of functional IGHD gene segments in horses in IgM and IgG repertoires.

**Supplementary Figure 2.**
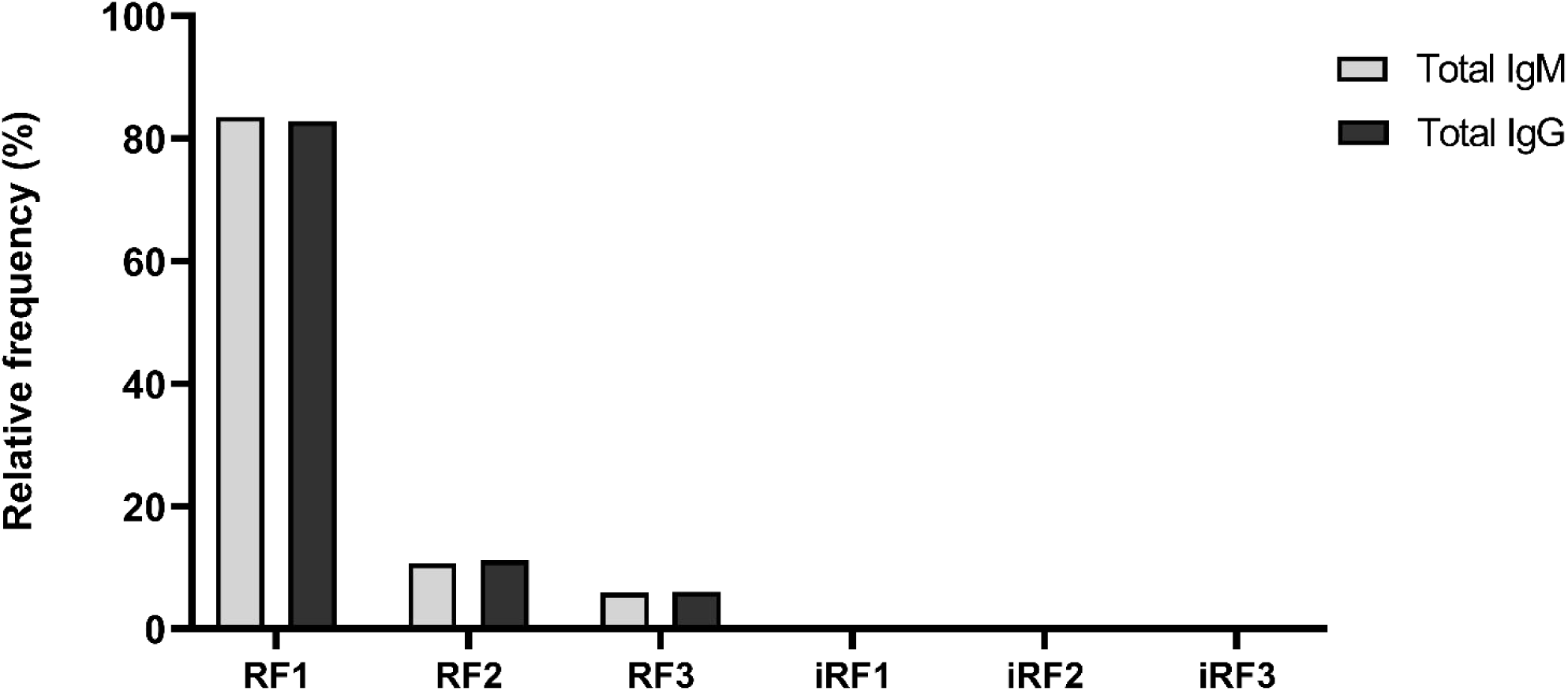
Reading Frame (RF) preference for IGHD gene segment in non-immunized horses. Relative frequency of the six possible open reading frames for the IGHD gene segment. RF1, RF2, and RF3 are generated by deletion, while iRF1, iRF2, and iRF3 are generated by inversion.

